# The salmon-peloton: hydraulic habitat shifts of adult Atlantic salmon (*Salmo salar*) due to behaviour thermoregulation

**DOI:** 10.1101/2021.04.19.440497

**Authors:** Antóin M. O’Sullivan, Tommi Linnansaari, Jaime Leavitt, Kurt M. Samways, Barret L. Kurylyk, R. Allen Curry

**Author notes:** corresponding author: Antóin M. O’Sullivan. **Conflict of Interest Statement** The author’s declare no conflict of interest.

## Abstract

In recent decades there has been an increase in conservation and restoration projects targeting Atlantic salmon (*Salmo salar* – AS), as populations in eastern Canada decline. Missing however, is an understanding of thermo-hydraulic habitat use by adult AS during summer, and thus the actual benefits of altering in-river physical structures. Here, we illustrated how optical and thermal infrared (TIR) imagery acquired from a UAV can be used in concert with *in-situ* depth and velocity data to map adult AS and develop models of thermo-hydraulic habitats in the Miramichi River, New Brunswick. We found during optimal thermal conditions (< 19 °C) proximity to boulders and Froude numbers, a non-dimensional hydraulic metric, were key parameters that characterized adult AS habitat. However, during behavioural thermoregulation events (>19 °C), proximity to the cool thermal plume and Froude number, a non-dimensional hydraulic parameter, were critical controls on habitat use. We also observed AS formed a distinct geometric formation during behavioural thermoregulation events, and term this formation a ‘*thermal-peloton*’. The primary function of the peloton is undoubtedly to reduce thermally induced stressed; however, we conceptualize the geometry of the peloton attenuates hydraulic-drag, and reduces energetic expenditure of individuals practicing behavioural thermoregulation. These data provide an unrivaled viewpoint of thermo-hydraulic habitat selection by adult AS, and a blue print for restoration work. The use of UAV-based sensors has the potential to instigate a paradigm shift for river sciences. The age of applying hyper-resolution, remote sensing for river science and aquatic ecology is immensely exciting.

## 1 Introduction

Atlantic salmon (*Salmo salar*) are a charismatic fish species native to many rivers in eastern North America, Northern and Western Europe, and Russia (Webb *et al*., 2007), and support multi-billion dollar fisheries. The physical attributes that determine suitable Atlantic salmon (AS) habitat, such as spawning habitat, have been linked to depth and velocity (Malcom *et al*., 2008). Moir et al. (2002) found Froude numbers, a non-dimensional hydraulic parameter based on the relationship between depth, velocity, and gravity, were strong predictors of spawning habitats. These hydraulic conditions are thought to remove fine clasts during redd construction and to help oxygenate eggs (Gillies and Moir, 2015). A plethora of studies have revealed the hydraulic regimes that characterize juvenile habitats during both summer and winter (*e*.*g*. Heggenes *et al*., 1990; Linnansaari and Cunjak, 2013). Heggenes, (1996) proposed that juvenile habitat selection is a function of potential net energy gains related to the velocity regime: access to drifting food availability define gains while swimming energy expenditure define costs. For adult AS returning to, and migrating up, rivers deep pools are suggested to be critical habitats (Crisp, 1996), while boulders have been linked to protection for predation (*e*.*g*., Keenleyside, 1962; Gibson, 1993).

While river hydraulics exert critical control on aquatic habitats, temperature is an equally, if not more, important determinate of suitable habitat (Breau *et al*., 2007) particularly for salmonids and other cold-water ectotherms (*e*.*g*., Morash *et al*., 2020). During summer, if water temperatures exceed critical thermal thresholds, AS can reduce metabolic and individual physiologic stress through ‘*behavioural thermoregulation*’ (Reynolds *et al*., 1979). This behaviour leads AS to seeking thermal refuge in cooler areas; cooler areas that are often sourced from shaded tributaries (Ebersole *et al*., 2003), groundwater-dominated tributaries (Corey *et al*., 2020), and springs (Torgersen *et al*., 1999). Thermal refuges may have unique hydraulic characteristics in addition to being thermally distinct (Ritter *et al*., 2020). For instance, Wilbur *et al*. (2020) found large sea-run brook trout (*Salvelinus fontinalis*) sought thermal refuges that were cold, but also deep.

Advancements in remote sensing has led to an exponential growth in the capacity to quantify physical features within fluvial environments (Torgersen *et al*., 1999; Legleiter and Harrison, 2019). Techniques for mapping river bathymetry include active methods, such as topobathymetric LiDAR mapping, and passive methods, such as optically derived bathymetric mapping (Marcus and Fonstad, 2010). While these mapping techniques first occurred in the geomorphic realm, they are beginning to be applied in ecohydraulic studies from reach (Lane *et al*., 2020) to catchment scales (O’Sullivan *et al*., 2020). As mapping techniques have evolved, concurrent advances in unmanned aerial vehicle (UAV) technology enable the mapping of Earth structures at unprecedented spatial and temporal scales. In the context of this study, recent research has illustrated the utility of bathymetric stream mapping by combing UAV obtained data and structure-form-motion techniques (Lane *et al*., 2020), as well as longitudinal stream temperature mapping using UAV-mounted thermal infrared (TIR) sensors (Casas-Mulet *et al*., 2020).

In recent decades there has been an increase in conservation and restoration projects targeting AS as populations in eastern Canada decline. These projects are often multifaceted including the restoration of not just hydraulic habitats (Lacey and Miller, 2004), but also thermal habitats (Justice *et al*., 2017). River hydraulic characteristics define adult AS habitat, and in summer, as river temperatures increase, temperature will affect habitat availability and selection – where AS select for thermal refuge to offset physiological stress (Frechette *et al*., 2018; Lennox *et al*., 2018). The occurrence of thermal refuges is a function of landscape characteristics and riverscape hydraulics (O’Sullivan *et al*., 2020; Ritter *et al*., 2020), and it appears both adult and juvenile AS can locate them (Frechette *et al*., 2018; Corey *et al*., 2020). These refuges are often known within local communities and by river managers who act to protect AS during predicted behavioural thermoregulation events (e.g., DFO, 2012). More recently, managers have noted that thermal habitats in summer may be augmented (Kurylyk *et al*., 2015; Ouellette *et al*., 2020) and have begun to alter instream habitats to create “cold water pools” for example (N. Wilbur, Regional Director Atlantic Salmon Federation, pers. comm.). Missing however is an understanding of thermo-hydraulic habitat use by adult AS during summer, and winter, and thus the actual benefits of altering in-river physical structures, *e*.*g*., the created hydraulic conditions, temporal persistent of created structure, and implications for the non-adult AS components of the ecosystem.

We use remote sensing techniques to passively quantify hydraulic preferences for adult AS before and during a thermal aggregation event on the Miramichi River, New Brunswick. We use TIR and red-green-blue (RGB) sensors mounted to an airborne UAV as well as *in-situ* depth and velocity data collected from an acoustic Doppler current profiler (ADCP) and RTK dGPS processed using a RF model approach. The AS behaviour was inferred from RGB data (e.g., Andrews at al. 2020) to develop movement patterns and habitat preference indices for each thermal condition (e.g., Mocq et al 2018; Wilbur *et al*., 2020). Our goal is to demonstrate the hydraulic conditions of preferred habitats for adult AS during a summer, behavioural thermoregulation event. In doing so, we hope to provide guidance for protecting and where necessary, creating adult AS habitat during the extreme temperatures of summer.

## 2 Methods

### 2.1 Study area

The study area is a 2.2 ha reach that includes the confluence of an unnamed tributary and the main stem of the Southwest Miramichi River, a northern temperate catchment in New Brunswick, Canada (Figure. 1). The Southwest Miramichi drains ∼ 7,700 km, and during summer is hydrologically characterised by low flows and warm temperatures (Caissie *et al*., 2007). Average August discharge from 1961-2018 = 74.9 m^3^·s^-1^, with average minimum discharge values for the same period = 19.5 m^3^·s^-1^ (Environment and Climate Change Canada (ECCC) gauge station: 01BO001). In the year of this study (August and early September 2020), the river experienced extreme low flows with an August minimum discharge = 8.6 m ^3^·s^-1^, the lowest August discharge since 1961. The long-term average maximum, average mean and average minimum air temperatures in August = 24.4 °C, 12.3 °C, and 18.4 °C, respectively (from 1943-2019; ECCC: 8100989). During these low flow, high air temperature situations, river water temperatures can exceed 30 °C (Breau *et al*., 2011; Corey *et al*., 2020). The 2 ha study site is located adjacent to a coldwater tributary that generates a coldwater plume in the main river, and adjacent to a private fishing club.

**Figure 1.**
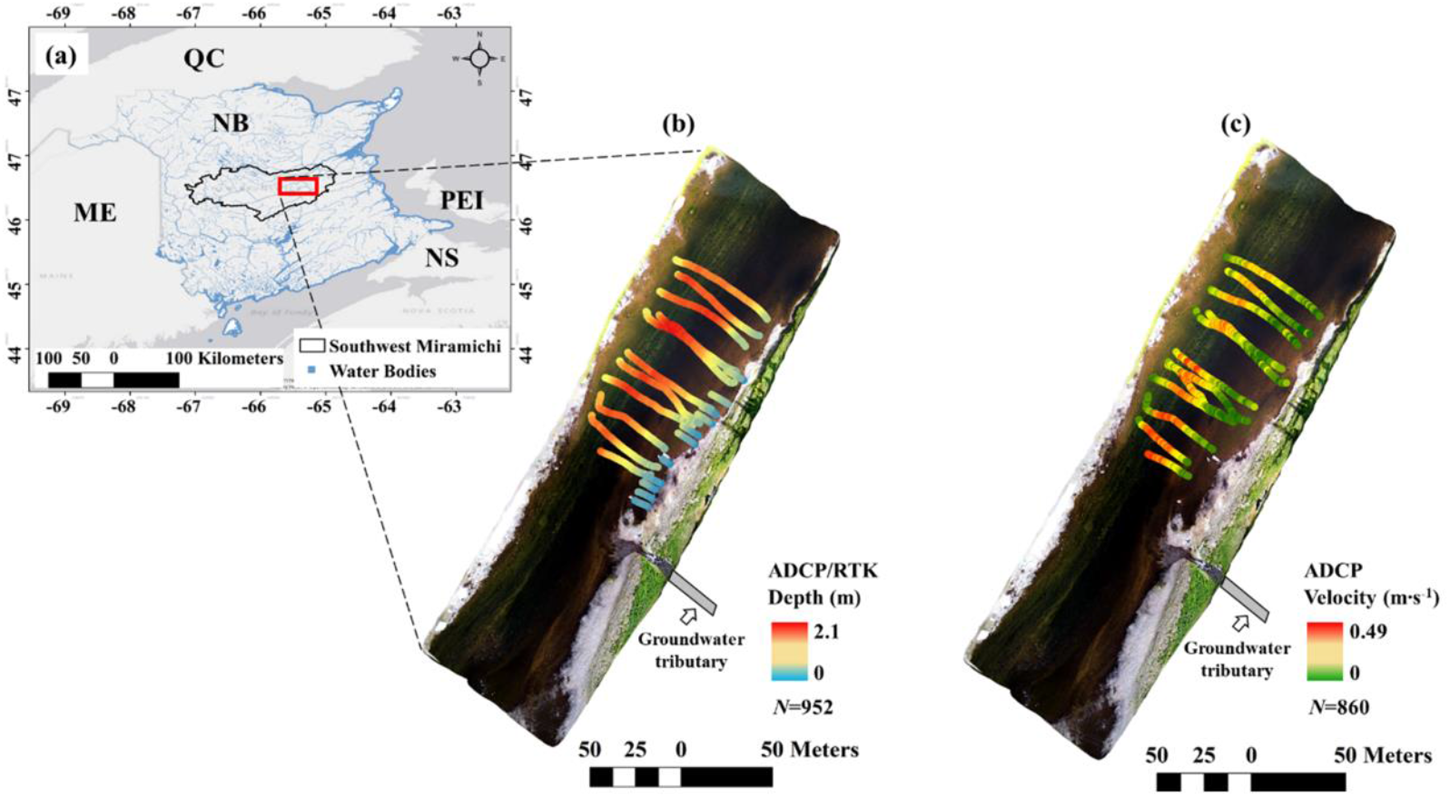
The geographic location of our study area located on the Southwest Miramichi, a tributary of the Miramichi River, New Brunswick (NB) denoted by the red polygon (a). Depth data were acquired via ADCP and RTK dGPS (b) and velocity data were collected via ADCP (c).

### 2.2 Workflow

The study approach followed four steps (Figure. 2). Step 1 involved the collection of UAV-based TIR data to delimit the spatial extent of the thermal plume, RGB optical imagery to observe AS aggregations and develop a suite of passive bathymetry models, and depth and velocity calibration data using a remote-controlled acoustic Doppler current profiler (ADCP), and Real-Time Kinematic and Differential GPS (RTK dGPS). Step 2 involved image and calibration data processing. In Step 3, hydraulic models were developed for the image datasets. Finally, in Step 4, preference curves were developed to investigate changes in thermo-hydraulic habitat selection before (optimal thermal conditions – see below) and during behavioural thermoregulation.

**Figure 2.**
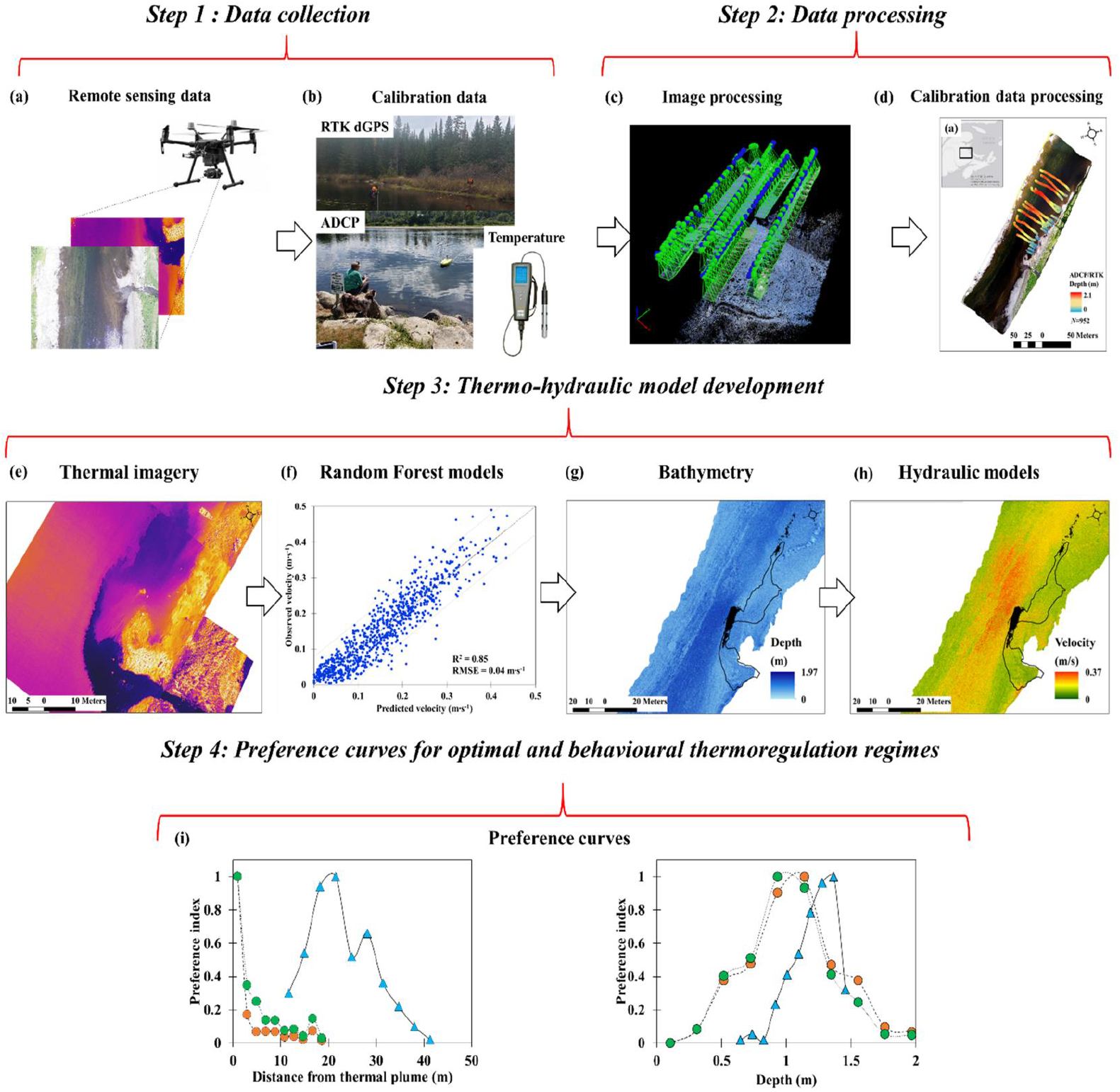
A workflow illustrating the steps to assess thermo-hydraulic conditions preferred by adult Atlanitc salmon for optimal and behavioural thermoregulation conditions.

### 2.3 Data collection

Adult AS often occupy well defined and pools during summer in the Southwest Miramichi (DFO, 2012). Our sample times were chosen when the local fishing camp guides observed behavioural thermoregulation and optimal thermal events, at which time we were alerted and conducted our data collection.

#### 2.3.1 Remote sensing imagery and field data

##### Remote sensing imagery

Remotely sensed imagery was collected from a DJI, Zenmuse XT2 thermal camera (19 mm) mounted to a DJI M210 RTK. The Zemmuse XT2 sensor is a dual FLIR Tau 2 thermal camera (7.5 – 13.5 µm) with a 4K RGB camera. Images for the behavioural thermoregulation conditions were collected at 10:30 August 24, 2020 and at 14:54 September 11, 2020. We defined ‘optimal thermal conditions’ as instances when AS are not observed to be behaviourally thermoregulating, and data for optimal thermal conditions were collected at 13:46 August 28, 2020. All days for image collection were cloudless, and water temperature of the main river was 19.1, 17.7 and 20.2 °C during August, 24, 28, and September 11, respectively (see Table 1). The tributary, and the apparent thermal refuge in the main river, was 14.3, 14.2, and 14.2 °C on each sampling day (Table 1).

**Table 1.** Sensor and flight information and hydroclimatic conditions for each mapping episode.

| Date/ Time | 8/24/2020; 10:30 | 8/28/2020; 13:46 | 9/11/2020; 14:54 |
| --- | --- | --- | --- |
| Atlantic Salmon behaviour | Behavioural thermoregulation | Optimal thermal conditions | Behavioural thermoregulation |
| Air temperature ($^{\circ}\text{C}$) | 25.4 | 21 | 18.1 |
| Main stem temperature ($^{\circ}\text{C}$) | 19.1 | 17.7 | 20.2 |
| Tributary temperature ($^{\circ}\text{C}$) | 14.3 | 14.2 | 14.2 |
| Water level (m)<br>(ECCC gauge station: 01BO001) | 0.37 | 0.44 | 0.37 |

Images were acquired from an altitude of 100 m with 60 % overlap. TIR imagery collected from uncooled thermal sensors, such as the FLIR Tau 2, can display a relatively large amount of thermal drift (see Dugdale *et al*., 2019), which can be corrected with calibration data from in-stream thermographs. However, the imagery was not calibrated in this study as the primary focus was to investigate changes in hydraulic preferences between optimal and behavioural thermoregulation conditions, *i*.*e*., the TIR images were used to delineate the thermal plume and demonstrate that AS were practicing behavioural thermoregulation. Point temperature measurements were taken for the air, main stem and tributary, using a YSI Sonde (± 0.3 °C; Table 1).

Both the RGB and TIR imagery were processed in Pix4DMapper © using the ‘3D model’ module. The image resolution was 3 cm for the RGB imagery and 7 cm for the TIR images. We resampled the RGB imagery to 5 cm resolution prior to development of optical derived depth models to reduce noise effects of image speckling (*e*.*g*., Jordan and Fonstad, 2005).

##### Field data

Depth and velocity data were synchronously collected on August 24 using an ADCP – ARCboat (HR Wallingford) sampled at 3 MHz. ADCP data were filtered (depths < 0.3 m removed) to remove potentially erroneous measurements (Guerro and Lamberti, 2011). Shallow depth measurements were collected using an RTK dGPS on August 24. In total *N* = 952 depth data points (*N* = 860 ADCP; *N =* 92 RTK GPS), and *N* = 860 velocity data points were collected (Figure. 1). The average, minimum, and maximum measured depths were 1.07 m, 0.01 m 2.1 m (SD = 0.6 m), respectively, and the average, minimum, and maximum measured velocities were 0.15 m·s^-1^, 0 m·s^-1^, and 0.49 m·s^-1^, respectively (Figure. 1 S1). Water levels, taken at the ECCC gauge station: 01BO001, were relatively stable during the study period (Table 1), and the August 24 depth and velocity data were used to calibrate image-derived hydraulic models for each day.

The extent of the August 28^th^ image was shorter than the August 24^th^ and September 11^th^ images. However, this was outside of the range of observed habitat use in the image, and did not affect the overall study. The reduction in image extent lead to a decrease in field calibration data points for the August 28^th^ image, where depth samples were reduced to *N* = 771 and velocity samples were reduced to *N* = 744.

### 2.4 Hydraulic and thermal characteristics

Passive optical bathymetric mapping (*e*.*g*., Legleiter and Harrison, 2019) exploits the relationship between light attenuation in the water column and depth to produce a bathymetric map (Marcus and Fonstad, 2010). In this study, we used the high-resolution RGB imagery to construct optical bathymetric maps. The red band (635 nm) in our imagery was found to best characterize depth variability and was chosen for the depth mapping exercise. A transform is required to capture the exponential nature of light attenuation in the water column (Lyzenga, 1981):

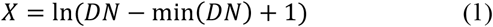

where *X* is an image-derived quantity, *DN* is the digital number of the band, and 1 is added to the expression as the natural log can only be calculated for positive values.

Machine-leaning algorithms are commonly used for optical depth mapping (*e*.*g*., Sagawa *et al*., 2020). We chose a Random Forest (RF) model and utilised the forest-based classification and regression tool in ArcGIS Pro (ESRI, 2020) calibrated with field depth measurements to predict depth from the *X* image from each observation. Similar to the depth mapping modelling approach, we used the forest-based classification and regression tool in ArcGIS Pro (ESRI, 2020) and the field velocity measurements to build velocity models for each observation.

Froude number (*Fr*), specifically the ratio of fluid (water) inertial to gravity forces, has shown strong correlations with AS spawning site selection (Moir *et al*., 2002). In this study we investigate the efficacy of *Fr* to explain habitat use by adult AS:

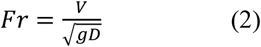

where *V* is the velocity of each pixel determined from the velocity RF model (m·s^-1^), *g* is the gravitational acceleration constant (9.81 m·s^-2^), and *d* is the depth of each pixel determined from the depth RF model (m). We modelled *Fr* using the forest based classification and regression tool in ArcGIS Pro (ESRI, 2020).

We manually delimited boulders and the thermal plume in ArcMap (ESRI, 2020) to mitigate against uncertainty.

### 2.5 Hydraulic habitat selection

#### Habitat preference indices

We manually delimited the spatial position of fish for all 3 dates in the RGB images. For optimal thermal conditions (August 28) *N* = 233 individuals were identified. It was not possible to demarcate individual fish for the dates of the apparent behavioural thermoregulation because the fish were aggregated closely together. Instead, a polygon was drawn around the aggregation for both August and September apparent behavioural thermoregulation events, along with any fish that could be individually identified proximal to the aggregation.

Following the delineation of AS, the polygons were overlain on the suite of depth, velocity and *Fr* maps, and values were extracted. To examine the influence of boulders and the thermal plume on habitat preferences, a proximity analyses was conducted in ArcMap using the ‘near (analysis)’ tool (ESRI, 2020), and the distance of fish from each feature was quantified.

Habitat preference indices are often used to describe habitat selection in aquatic studies (see Heggens, 1996). We calculated habitat preference indices by:

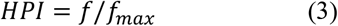

where *HPI* is the habitat preference index, *f* is the frequency individuals are observed to select for a certain habitat type, (temperature, depth, velocity, *Fr*, and distance from boulders), and *f*_*max*_ is the maximum observed frequency of a habitat type selected. Preference indices range from 0 (not selected) to 1 indicating preferred habitat.

## 3 Results

### 3.1 Adult Atlantic salmon behaviour

The apparent behavioural thermoregulation events were observed on August 24 and September 11 (Figure 3a and c). The optimal (non-event) behaviour is shown for August 28 (Figure 3b), and the coldwater plume in the main river on August is shown in Figure 3d.

**Figure 3.**
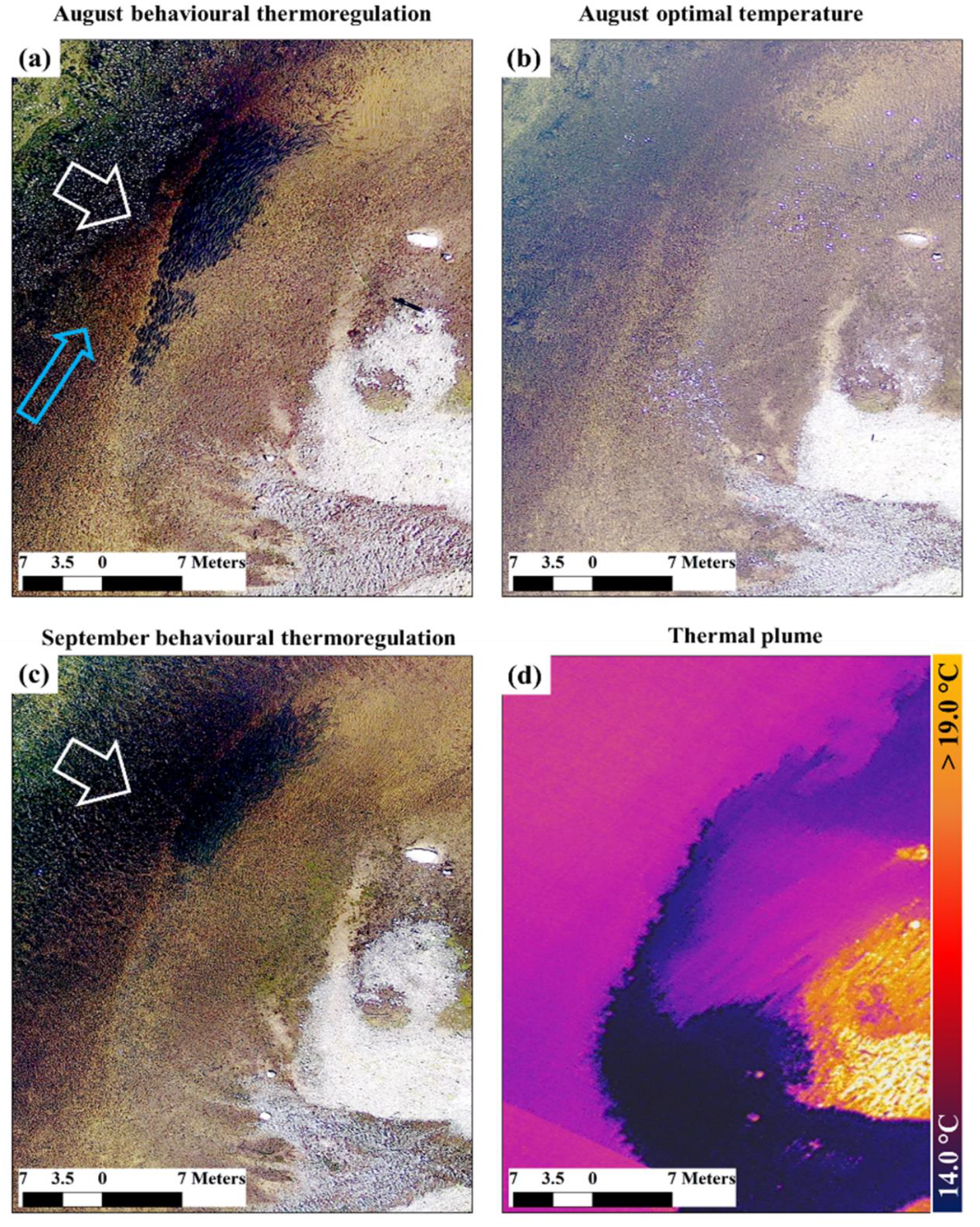
The behaviour of adult Atlantic salmon during two apparent, behavioural thermoregulation events on August 24 (a) and September 11 (c). Aggregations are demarcated with the white arrows, and flow direction by the blue arrow. During non-events, adult salmon are widely dispersed with no aggregations (b). The thermal plume, approximately 14 °C, created by the adjacent tributary entering the main river is shown in (d).

### 3.2 Hydraulic models

The RF generated depth models were relatively robust for each event, with *Adj. R*^*2*^ values ranging from 0.79 – 0.82 and *RMSE* between 0.26 and 0.28 m (see Table 2; Figure S2). Similarly, the velocity and *Fr* models showed relatively high, and consistent, levels of accuracy with *Adj. R*^*2*^ = 0.77 – 0.86 (*RMSE* = 0.04 – 0.05 m·s^-1^), and *Adj. R*^*2*^ = 0.86 (*RMSE* = 0.01), respectively (see Table 2; Figure S3). For both apparent behavioural thermoregulation events, fish were located within or close to the thermal plume, and avoided the thalweg, the section with the highest velocity and *Fr* values (Figure 4a-c, and 4g-i). During optimal thermal regime conditions, fish were observed opposite the plume, but still avoided the thalweg (Figure 4d-f).

**Table 2.** Optically derived bathymetric, and hydraulic model results for each event.

| Hydraulic parameter | August behavioural thermoregulation regime | August optimal thermal regime | September behavioural thermoregulation regime |
| --- | --- | --- | --- |
| Depth (m) | *Adj. R*<sup>2</sup> = 0.82 | *Adj. R*<sup>2</sup> = 0.79 | *Adj. R*<sup>2</sup> = 0.81 |
|  | *RMSE* = 0.26 | *RMSE* = 0.28 | *RMSE* = 0.26 |
|  | (*N* = 952) | (*N* = 771) | (*N* = 952) |
| Velocity (m·s<sup>-1</sup>) | *Adj. R*<sup>2</sup> = 0.77 | *Adj. R*<sup>2</sup> = 0.81 | *Adj. R*<sup>2</sup> = 0.86 |
|  | *RMSE* = 0.05 | *RMSE* = 0.05 | *RMSE* = 0.04 |
|  | (*N* = 860) | (*N* = 744) | (*N* = 860) |
| Froude | *Adj. R*<sup>2</sup> = 0.86 | *Adj. R*<sup>2</sup> = 0.85 | *Adj. R*<sup>2</sup> = 0.86 |
|  | *RMSE* = 0.01 | *RMSE* = 0.01 | *RMSE* = 0.01 |
|  | (*N* = 860) | (*N* = 744) | (*N* = 860) |

**Figure 4.**
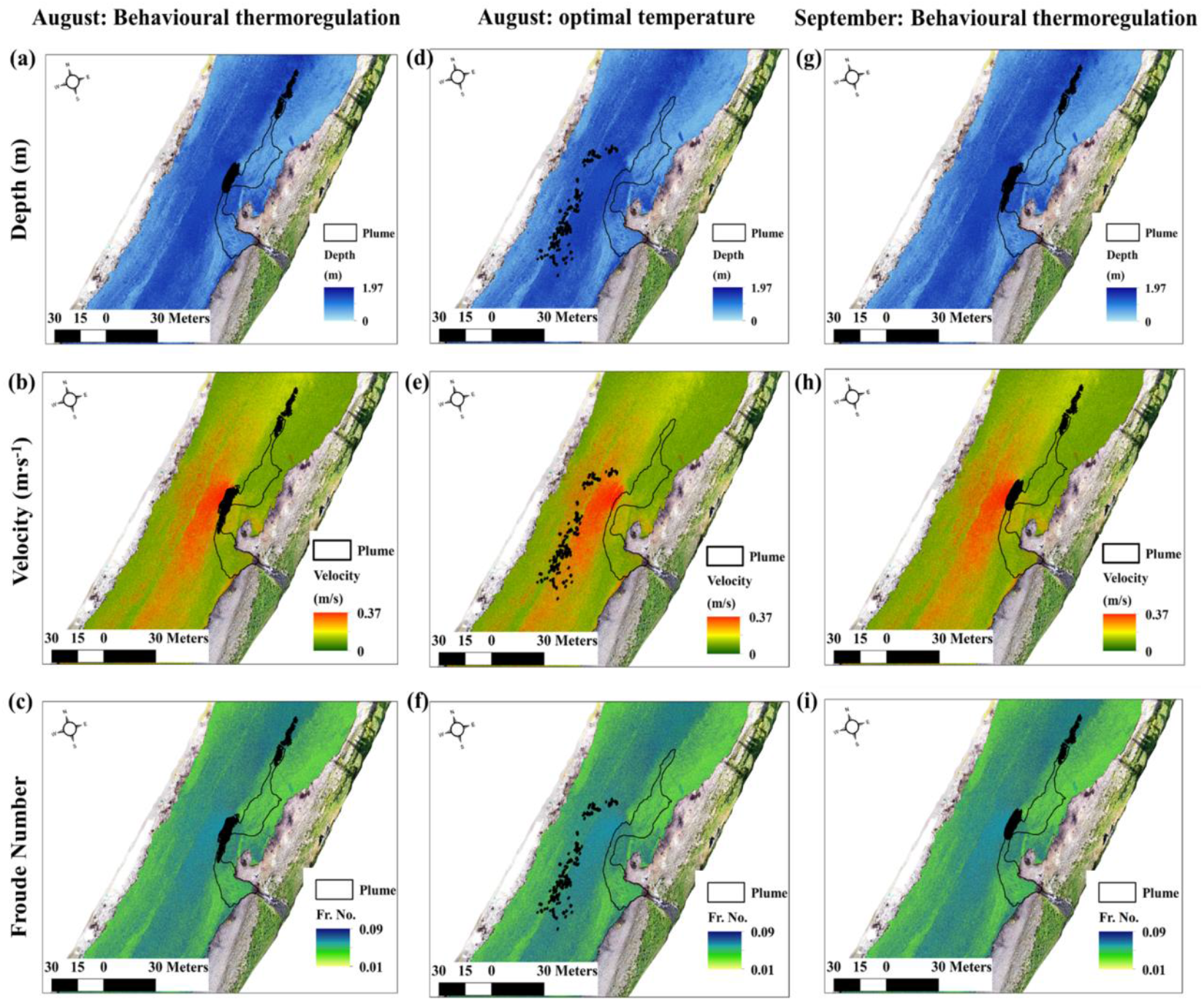
The position of adult Atlantic salmon (black features) during the apparent behavioural thermoregulation event in relation to the thermal plume (black polygon) on August 24 in relation to depth (a), velocity (b), and Froude number (c), and similarly for the optimal thermal conditions on August 28 (d, e, f), and the September 11 behavioural thermoregulation event (g, h, i).

### 3.3 Hydraulic preference curves

On the day of optimal thermal conditions (August 28) a total of *N* = 233 fish, occupying ∼ 14.8 m^2^, were identified in the image. On average each fish polygon for the optimal event had an areal extent = 0.06 m^2^, encompassing ∼ 120 pixels. Depth preferences for the optimal thermal condition (August 28), were observed to be ∼ 1.3 m, velocity preferences were ∼ 0.2 m·s^-1^, and *Fr* preferences were ∼ 0.07, while habitats located within < 1 m from boulders were noticeably preferred (Figure. 5). Distances ∼ 24 m from the thermal plume were preferred (Figure 5).

**Figure 5.**
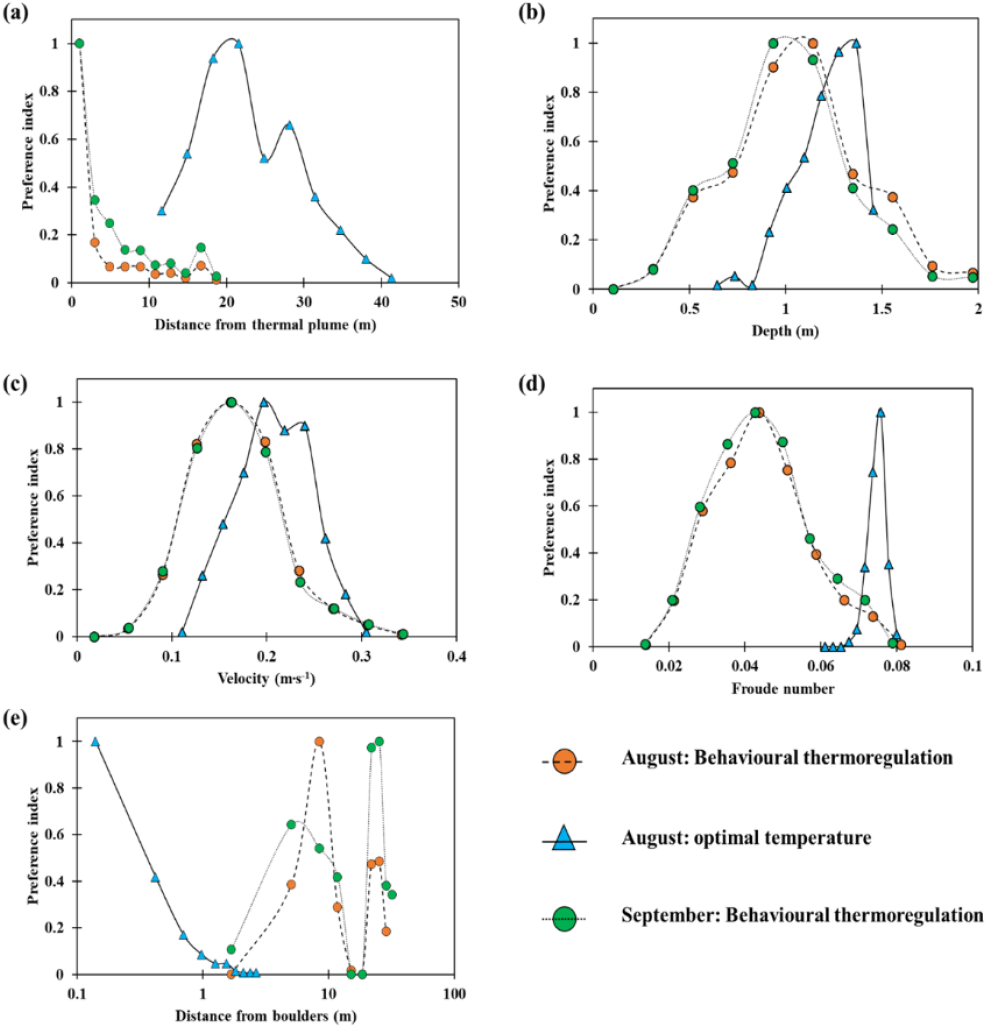
Habitat preference indices for optimal and behavioural thermoregulation conditions: distance from the thermal plume (a), depth (b), velocity (c), Froude number (d) and distance from boulders (e).

The polygon that delimited the August 24 behavioural thermoregulation event was ∼ 74.4 m^2^, and the areal extent of the September 11 behavioural thermoregulation event polygon was 68 m^2^ (see Figure 5). Preference indices for depth, velocity, *Fr*, and distance to thermal plume were similar for both behavioural thermoregulation events, and markedly different from optimal thermal conditions (Figure 5). Depth preferences for the behavioural thermoregulations were observed to be ∼ 1.1 m, velocity preferences were ∼ 0.16 m·s^-1^, and *Fr* preferences were ∼ 0.04. Distances within 1 m of the thermal plume were preferred (Figure 5).

## 4 Discussion

### 4.1 Summer thermo-hydraulic habitats of adult Atlantic salmon

Using remotely sensed data for river temperature (TIR) and depth (RGB), we mapped a change in summer habitat use for adult AS between optimal conditions and behavioural thermoregulation events. Our UAV images and ADCP data produced fine scale (5 cm) maps of adult locations, river thermo-hydraulics and in-stream features, *i*.*e*., boulders. From these data, we reveal that habitat preferences for optimal and behavioural thermoregulation events. The prior includes preferences for boulders, depth and velocity, while the latter details a preference for the coldwater plume and low *Fr* values.

Temperature preference for adult AS ∼ 16.5 – 17.5 °C (Stehfest *et al*., 2017). Under stressful thermal conditions, they will move amongst mesohabitats in a river, which is attributed to behavioural thermoregulation (Dugdale *et al*., 2016; Wilbur *et al*., 2020). Our fine scale observations demonstrate that movements can capitalize on coldwater plumes created by, for example, inflowing tributaries (Otter brook, Little Southwest Miramichi – Corey *et al*., 2020). At our study location, the thermoregulation behaviour occurred when river temperatures outside the plume reached 19°C. This is lower than behavioural thermaloreglation temperatures found by Shepard, (1995) in the Penobscot, Maine (USA), but is consistent with the findings of Frechette *et al*., (2018) in the Rivière Sainte-Marguerite Nord-Est, in Quebec, Canada.

Our fine scale observations suggests that high temperature events alter not only thermal habitat preferences but also hydraulic preferences. Adult AS do no feed in the summer as they migrate upstream to spawning areas triggered by fluxes in flow (Erkinaro *et al*., 1999); therefore, habitat selection is driven by energy conservation, and predation avoidance (Jones, 1952). Deep pools, boulders, and overhung banks are often selected (Crisp, 1996); but interestingly, we observed the deepest (< 1.9 m), and darkest, sections of the pool were not utilised during optimal thermal conditions (Figure 4). The deepest pool sections of our study area also had the highest velocities (> 0.3 m·s^-1^) and lacked boulders (Figure. 4). Instead, adult AS were observed near boulder fields (within 1 m), in areas with depths ∼ 1.3 m, and velocities ∼ 0.2 m·s^-1^ (Figure 5). Additionally, we observed adult AS selecting areas exposed to sunlight, but within close proximity to boulders (Figure S3). Adult AS showed a clear preference for *Fr =* 0.07 (Figure 5). We hypothesize that the selection of boulder habitats, and *Fr values* = 0.07 is multifaceted. While boulders do offer cover, they also create micro-hydraulic habitats, or provide ‘hydraulic-refuge’.

Boulder fields cause complex turbulence regimes and associated eddy structures that alter velocity regimes. Changes in hydraulics include deceleration at the front of boulders, recirculation immediately downstream of the boulders in a region termed the “near-wake” region where flows reverse and move back upstream, and finally deceleration close to the bed downstream of the near wake region in a region termed the “far-wake” region (Papanicolaou *et al*., 2012). Papanicolaou *et al*. (2012) found that velocity modulations due to the presence of boulders were observed downstream, up to a length of 3.5 times the boulders diameter. We hypothesize that adult AS capitalize on the energy efficiencies provided by boulders, utilising both near- and far-wake regions during optimal thermal conditions (see Figure S3). The boulders likely provide hydraulic refuge where the recirculating upstream currents significantly reduce energetic costs (Jonsson *et al*., 2007). A proximity to boulders by adult AS is a well-understood characteristic among anglers. Interestingly, it was revealing to observe the adults moving away from boulders as river temperatures rose to physiologically stressful levels.

Adult AS also moved into shallower and lower velocity waters during the thermally stressful events (Figure 4 and 5). Our results suggest that these stressful events push adults away from generally preferred energetic conservation and predator protection locations, e.g., boulders and deep water locations, to access the cooler water. It was apparent that the best hydraulic indicators of preferred habitat during this thermoregulation behaviour were both velocity and *Fr* (Figure 4), which averaged 0.15 m.s^-1^ and 0.04, respectively, and were and clearly less than values during optimal thermal conditions. Low velocity and *Fr* values indicate sub-critical flow, thereby areas where less energy would be expended. This illustrates that thermal refuges for adult AS are defined by temperature, but may also be identified by depth and velocity characteristics, best represented by *Fr* number.

The high summer temperature events also pushed the AS into aggregations similar to the behaviour of juveniles during these events (Breau *et al*., 2007, Corey *et al*., 2020). The geometry created by aggregating adult AS displayed obclavate shapes (Figure 4 and 6a and b). The primary driver is undoubtedly to provide access to the thermal plume (Frechette *et al*., 2020); however, the geometry of the aggregation is similar to that utilised by cyclists to reduce aerodynamic-drag, a formation known as a ‘peloton’. Recent work by Blocken *et al*., (2018) found aerodynamic drag was reduced by 90 – 95 % in the mid-rear of the peloton when compared to an isolated cyclist. This behaviour is also commonly used in schooling fishes to dampen hydraulic-drag (*e*.*g*., Weihs, 1975; Daghooghi and Borazjani, 2015). We present a concept we term a ‘*thermal-peloton*’. We hypothesize that the observed thermal-peloton used by adult AS (Figure 6a) reduces thermal stress, whilst also favourably modulates hydrodynamics. We predict that the geometry of the thermal peloton attenuates hydraulic drag, thereby reducing energetic expenditure for the group, but with potential costs for some individuals, *e*.*g*., the front of the peloton (see review by Trenchard and Perc, 2016). Our observations are static, thus more studies will be required to establish if a salmon peloton functions like the dynamic, human cyclist peloton where social interactions lead to individuals moving among the high and low energy expenditure positions (Blocken *et al*., 2018).

**Figure 6.**
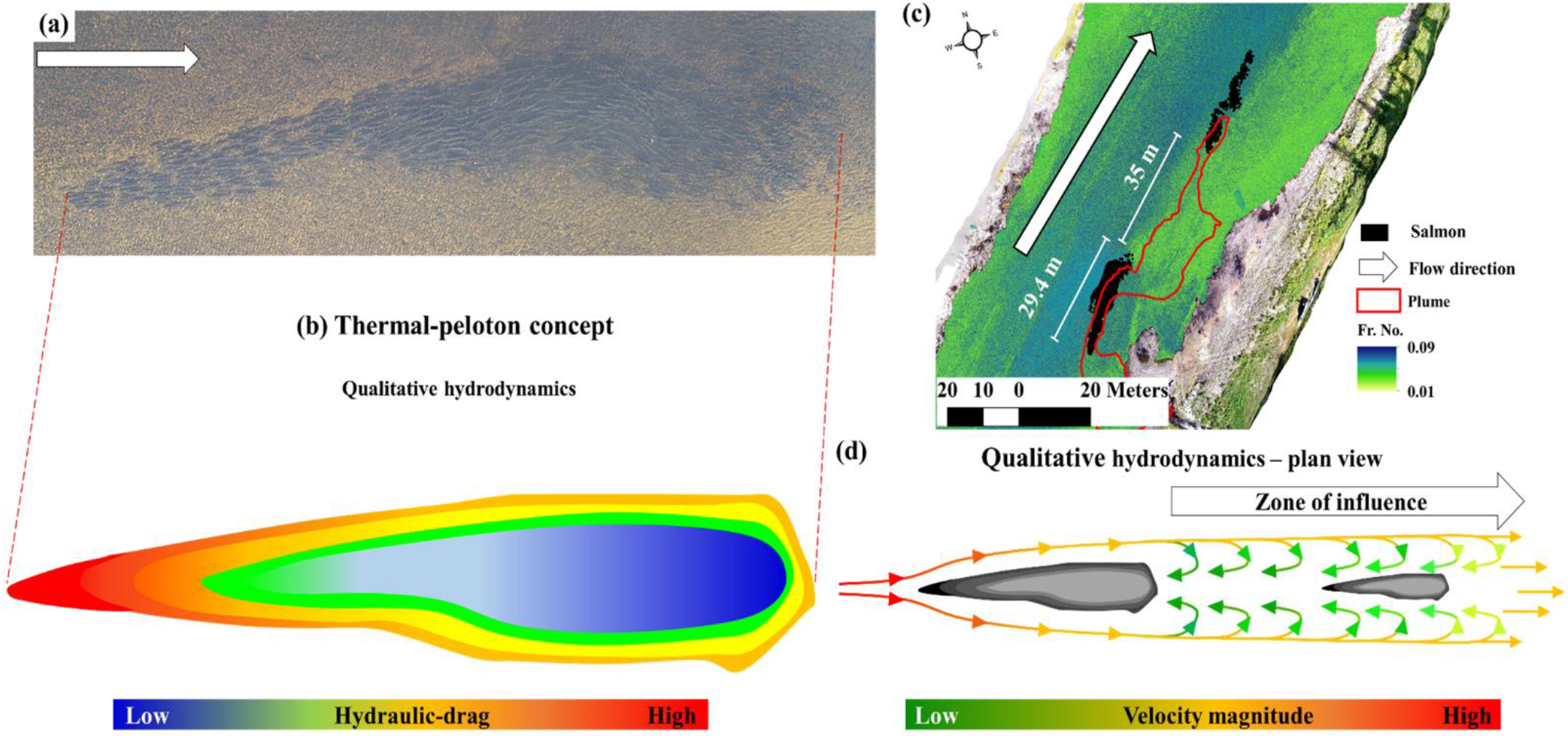
A behavioural thermoregulation event for adult Atlantic salmon and the concept of a ‘thermal peloton” (see text). Two aggregations of individuals existed; the upstream the upstream aggregation was larger (a) and (b) represents a schematic prediction of the group’s hydraulic drag. The two aggregations in relation to the thermal plume (c), and (d) represents a schematic prediction of the velocity associate with the grou

We observed two groups of adult AS during thermally stressfully conditions (Figure 4). The downstream group also appear to selected the deepest remaining area of the plume with lower *Fr* values ∼ 0.04 (Figure 5 and 6c). It is worth noting that TIR only measures the surface of the water, and mixing processes likely extend the plume beyond the observed surface extent (Handcock *et al*., 2012). The hydraulic conditions do however suggest hydrodynamic benefits to the downstream group due to the larger upstream group, which acts as a hydraulic structure that modulates hydraulic regimes around and behind the upstream group. Extrapolating the findings of Papanicolaou *et al*., (2012), the ∼ 29 m long upstream group could generate a zone of hydraulic influence that extends past the downstream group, *i*.*e*., assuming a zone of influence 3.5 time the length of the group (Figure 6d). As such, the upstream group most likely provides a bio-derived hydraulic refuge for the smaller downstream group.

### 4.2 Utility of hyper-resolution sensors for ecological studies

The fine-scale resolution of the thermal-hydraulics of the AS behaviour are extraordinary, but remotely sensed images and machine learning models are not without caveats. Passive depth mapping is underpinned by the physics of light attenuation in water *(e*.*g*., O’Sullivan *et al*., 2020), thus both colour and surface smoothness may interfere with interpretations. TIR temperatures only measure the surface of the water and in-river mixing processes likely shape the plume beyond the observed 2-dimensional, surface extent (*e*.*g*., Handcock *et al*., 2012); this may be why some individual AS appear “outside” the plume during the high temperature events (upper peloton). Our hydraulic variable maps were derived from a machine learning approach (Random Forest) and while emerging as a very powerful tools in the natural sciences, the approach has limitations, particularly machine learning models are poor extrapolators (*e*.*g*., Tyralis *et al*., 2019). The hydraulic conditions we report are consistent with our collective, years of experience on the Miramichi River (Burge, 2005); we and others will continue to explore the power and limitations of machine leaning models in river sciences (e.g., Tyralis *et al*., 2019; Zhu *et al*., 2020)

Remote sensing has revolutionized global and regional scale studies across myriad research fields, such as ecology (Torgersen *et al*., 1999), and geomorphology (Legleiter and Harrison, 2019). In this study, we illustrated how optical imagery acquired from a UAV can be used in concert with *in-situ* depth and velocity data to passively map adult AS and develop models of hydraulic habitats, and how paired thermal infrared imagery can be applied to detail thermal heterogeneity. These data provide an unrivaled viewpoint of behaviour and its link to hydraulic conditions, and consequently, open doors for rapid advances in river ecology and provides a blueprint for habitat restoration work. While current habitat restoration is mostly based on coarse scale interpretations of river hydraulics and engineering principles (see recent review by Adeva-Bustos *et al*., 2019), our findings indicate that the preferred habitat for adult AS during high temperature events is best described by fine-scale, *Fr*. Adults also endure stress related to habitat restrictions during winter, and it is likely that the hydraulics under and around river ice used by AS will be best described by hydraulic variables such as *Fr* numbers that are already linked to other habitat choices, *e*.*g*., redd sites (Moir et al. 2002). We have demonstrated that such advances in ecology and hydraulic engineering using UAV-based sensors has the potential to instigate a paradigm shift for river sciences (Dugdale *et al*., 2019; Casa-mullet *et al*., 2020). The age of applying hyper-resolution, remote sensing for river science and aquatic ecology is immensely exciting.

## Data availability statement

The data in this study detail a critical thermal refuge that could be easily exploited if its location was revealed. The data generated within are georeferenced. In the interest of conservation, the data are being held by the lead author. Any queries regarding these data can be addressed to the lead author.

## Acknowledgements

First, the authors would like to thank Lord Pisces. The authors wish to thank the Collaboration for Atlantic Salmon, J.D. Irving, Limited, Cooke Aquaculture, NSERC (RGPIN-2018-06015 – RAC) and New Brunswick’s Wildlife Trust Fund. The work also received partial financial contributions from the Fisheries and Oceans Canada / Ce Project fut partiellement appuyé parune contribution financière de Pêches et Océans Canada, The Atlantic Salmon Conservation Foundation, Mirmaichi Salmon Association, and New Brunswick’s Innovation Foundation. We thank K. Haralampides, and D. Connor for assistance collecting ADCP data. We also thank an unnamed fishing club on the Southwest Miramichi for site access.

