## Supplemental information for "The salmon-peloton: hydraulic habitat shifts of adult Atlantic salmon (*Salmo salar*) due to behaviour thermoregulation"

***Running title***: The salmon-peloton: thermo-hydraulic habitats of adult Atlantic salmon

Antóin M. O'Sullivan[1,2*], Tommi Linnansaari[1,2,3], Jaime Leavitt[1,4], Kurt M. Samways[1,5], Barret L. Kurylyk[6], and R. Allen Curry[1,2,3]

[1] Canadian Rivers Institute

[2] FOREM, University of New Brunswick (Fredericton)

[3]Biology, University of New Brunswick (Fredericton)

[4]Civil Engineering, University of New Brunswick (Fredericton)

[5]Biological Sciences, University of New Brunswick (Saint John)

[6]Civil and Resource Engineering, Dalhousie University (Halifax)

**Contents of this file**

Figure S1 to S3

**Introduction**

This supporting information provides additional details to those in the main paper.

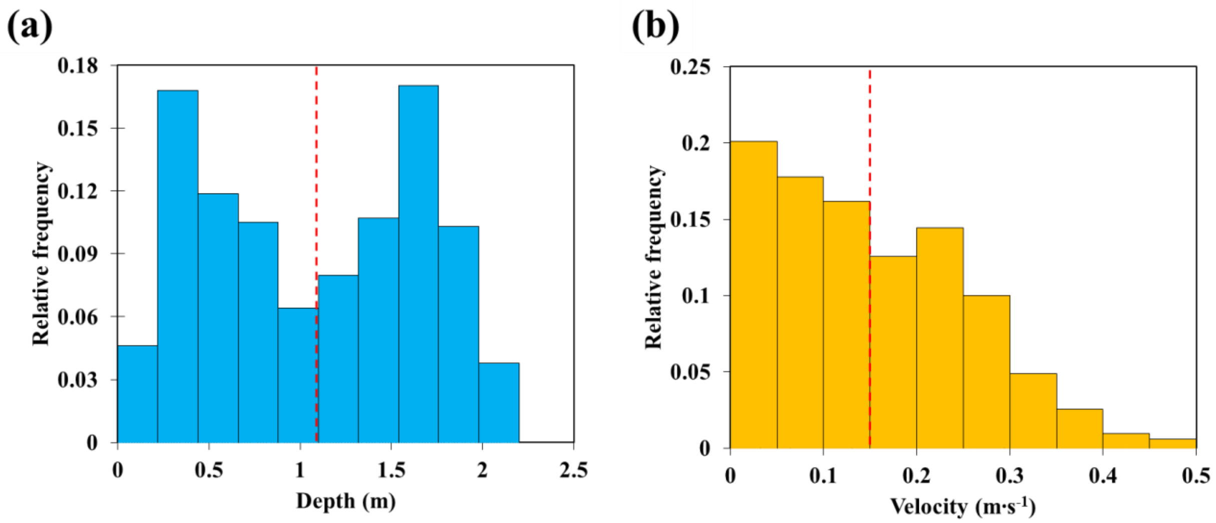


**Figure S1** The relative frequency distribution of depth (a – *N* = 952) and velocity data (b – *N* = 860) collected via ADCP and RTK dGPS during August 24 2020.

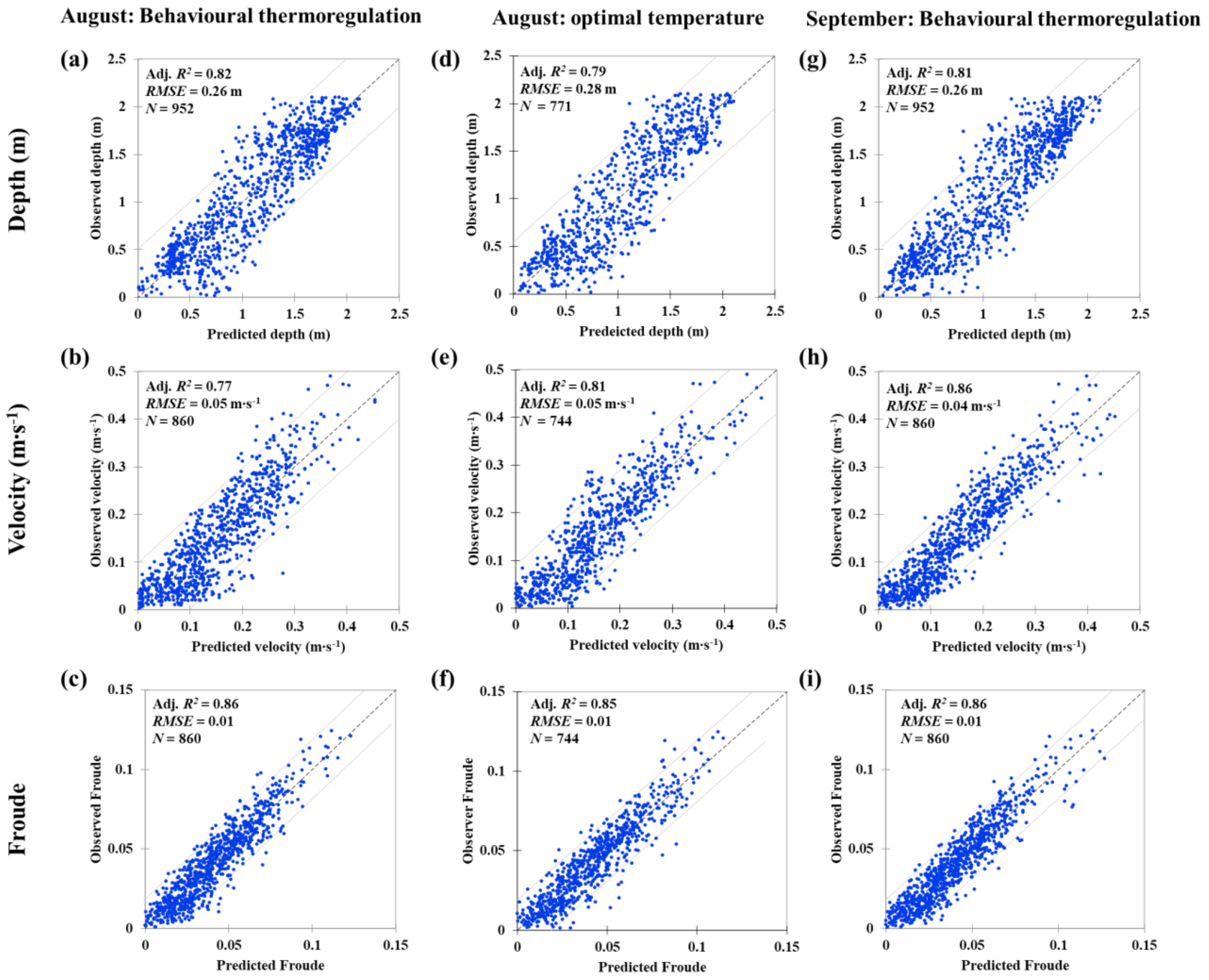


**Figure S2** Model fits for the August behavioural thermoregulation depth (a), velocity (b) and Froude (c) models. Depth, velocity and Froude model fits for optimal temperature conditions, and September behavioural thermoregulation conditions are presented in (d), (e), (f), (g), (h), and (i), respectively.

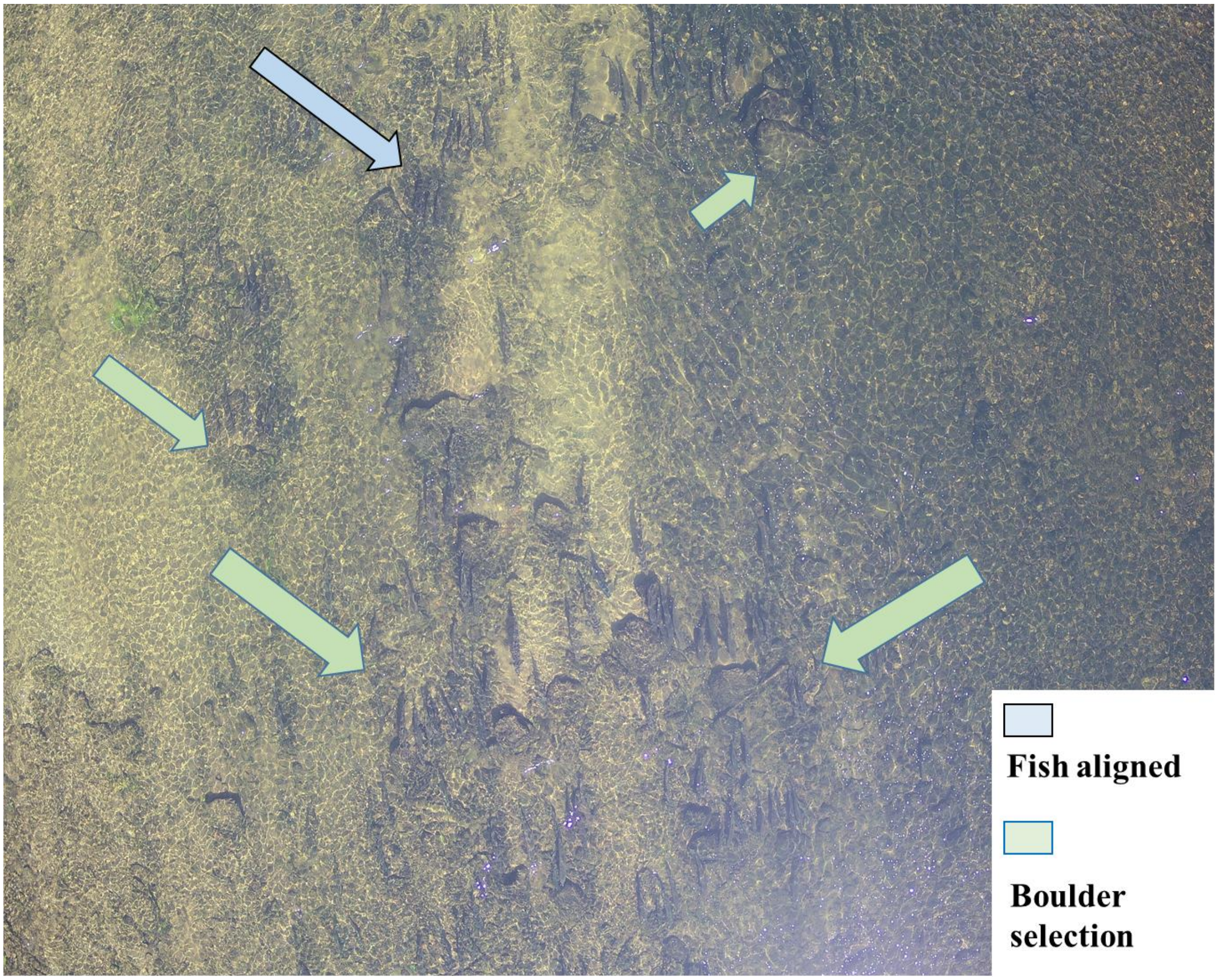


**Figure S3** An aerial image captured from an optical sensor revealing adult Atlantic salmon habitat use where salmon are absent from areas without boulders, and clustered in areas with boulders (olive arrow). Adult salmon are also observed aligned in a down river direction (blue arrow).
